# SMASH: Scalable Method for Analyzing Spatial Heterogeneity of genes in spatial transcriptomics data

**DOI:** 10.1101/2023.03.23.533980

**Authors:** Souvik Seal, Benjamin G. Bitler, Debashis Ghosh

## Abstract

In high-throughput spatial transcriptomics (ST) studies, it is of great interest to identify the genes whose level of expression in a tissue covaries with the spatial location of cells/spots. Such genes, also known as spatially variable genes (SVGs), can be crucial to the biological understanding of both structural and functional characteristics of complex tissues. Existing methods for detecting SVGs either suffer from huge computational demand or significantly lack statistical power. We propose a non-parametric method termed SMASH that achieves a balance between the above two problems. We compare SMASH with other existing methods in varying simulation scenarios demonstrating its superior statistical power and robustness. We apply the method to four ST datasets from different platforms revealing interesting biological insights.

## Background

Spatial transcriptomics (ST) performs high-throughput measurement of transcriptomes in complex biological tissues at single-cell or subcellular resolution, preserving spatial information [1, 2, 3, 4, 5, 6, 7, 8, 9, 10]. In the past decade, the rapid development of ST technologies has facilitated exciting discoveries in different domains, including neuroscience [11, 12, 13] and cancer research [14, 15, 16]. Early ST technologies, such as smFISH [17], seqFISH [18] and MERFISH [19], operate at relatively low spatial resolution, whereas recent technologies, such as Slide-seq [20], Slide-seq V2 [21], 10X Visium [22] and HDST [23], have enabled transcriptome-wide profiling at a much finer spatial resolution on multiple thousands of locations. Because of such huge spatial profiles, deriving biological insights from these datasets not only poses a plethora of statistical challenges but also demands maximal computational efficiency [24].

A critical step in the analysis of ST datasets is to identify the genes whose level of expression co-varies with the spatial locations across the tissue. These genes, often referred to as spatially variable genes (SVGs), can be used in downstream analyses, such as identifying potential markers for biological processes and defining areas in the tissue that dictate cellular differentiation and function [25, 26, 27, 28, 29]. For example, Wang et al. (2020) [30] analyzed an ST dataset on the tumor microenvironment (TME) of three tissue sections from a prostate cancer subject [31]. In every tissue section, a unique set of spatially variable metabolic genes were identified, which they argued, can be used to guide targeted tissue-specific therapy. A simplistic approach for detecting SVGs could be to identify spatially located layers or cell types (if any) based on either a priori biological knowledge or using popular softwares, such as RCTD [32] and Seurat [33], with the transcriptional profiles, and then check which genes are enriched more in a particular spatial layer or cell type. However, such an approach would only work if the cell types are spatially well-separated, and always be sensitive to the quality of the cell type-identification step [34]. In recent years, more sophisticated methods have been developed to identify SVGs, a systematic overview of some of which can be found in Li et al. (2021) [35]. The methods can be broadly classified into two types: a) methods based on statistical modeling and b) methods based on graphical modeling. The methods of type (a) are based on parametric or non-parametric statistical model assumptions, whereas the methods of type (b) are usually model-free and based on graphical networks, constructed directly from the spatial locations or using simplified spatial grids. Some of the notable methods of type (a) include Trendsceek [36], SpatialDE [34], SPARK [37], and SPARK-X [38]. Some of the methods of type (b) are HMRF [39], MERINGUE [40], SpaGCN [41] and SpaGene [42]. Being model-based, type (a) methods allow adjusting for covariates such as cell type, whereas type (b) methods claim to be more flexible in the sense of being model-free.

The statistical power of both types of methods greatly varies based on gene expression patterns and the spatial structure of an ST dataset. The methods encounter different levels of computational complexity based on two quantities, *N* and *K*, corresponding to the numbers of cells/spots and genes, respectively. SpatialDE [34] is one of the earliest methods of type (a). It employs a Gaussian process (GP) regression model [43] with kernel-based covariance matrices [44] of multiple types, such as linear, Gaussian, and cosine, computed using the distance between the spatial coordinates of the cells. The model decomposes the total variability of a gene expression into two components, spatial and error variance. A significantly large value of the spatial variance would imply that the gene is spatially variable. Borrowing an efficient estimation algorithm from the statistical genetics literature [45], SpatialDE manages to estimate the variance components with a reasonable degree of computational efficiency, requiring *O*(*N* ^3^ + *N*^2^*K*) floating point operations (FLOPS). A newer method named SPARK [37] extends the framework of SpatialDE by considering a generalized linear spatial model (GLSM) [46] with a Poisson distribution, arguing to be better suited for modeling the raw count data usually obtained from most of the ST platforms. However, the penalized quasi-likelihood (PQL) approach [47] used for parameter estimation in SPARK is extremely computationally demanding with a complexity of *O*(*N* ^3^*K*), making it unusable for a transcriptome-wide analysis when *N* is moderately large (*N >* 3, 000). To this end, a non-parametric highly scalable method named SPARK-X [38] has been recently developed requiring just linear complexity w.r.t. *N*. It is based on the robust covariance testing framework [48] that compares the linear kernel-based covariance matrices of the gene expression and the spatial coordinates. However, using a linear kernel makes SPARK-X equivalent to fitting a multiple linear regression model [49] with the gene expression as the dependent variable and the spatial coordinates (or, some transformation of these) as the predictors and testing if the fixed effect coefficients differ from zero. Thus, it is only capable of detecting spatial dependencies or patterns that manifest linearly in the mean or expected value of the gene expression, also known as first-order dependencies, and drastically loses power in complex scenarios as to be shown later. Zhu et al. (2021) [38] have partially acknowledged this issue with their main focus being computational scalability.

On the other side, a popular method of type (b), MERINGUE [40] considers spatial autocorrelation and cross-correlation based on spatial neighborhood graphs to identify SVGs. Improving hugely on the complexity of MERINGUE, another model-free method named SpaGene [42] has been recently developed. It constructs a spatial network between cells/spots using the *k*-nearest neighbors approach, and then for each gene, extracts the subnetwork whose nodes have high gene expression. Then, it compares the observed degree distribution of the subnetwork to a distribution from a fully connected network using the earth mover’s distance [50]. It considers a permutation test [51] to obtain the *p*-value for every gene. SpaGene is highly comparable to SPARK-X w.r.t. computational complexity and thus applicable to ST datasets with large *N*. However, the method is harder to interpret than the methods of type (a), can not readily accommodate additional covariates, and also lacks power in various scenarios (see Simulations section).

We propose a non-parametric method, named SMASH, which achieves superior statistical power than both SPARK-X and SpaGene, while remaining computationally tractable. It augments the idea of SPARK-X in its use of the robust covariance testing framework [48] coupled with more general kernel-based spatial covariance matrices. With a computational complexity quadratic in *N*, SMASH sacrifices some degree of computational efficiency in favor of significantly higher detection power than both SPARK-X and SpaGene. However, it is worth highlighting that SMASH is notably faster than other type (a) methods, such as SpatialDE and SPARK, and can thus be thought of as a balanced alternative, fusing high detection power with a moderate degree of scalability. In varying simulation scenarios, we demonstrate that SMASH achieves highly consistent and superior performance as compared to the methods SPARK-X and SpaGene. Finally, our analysis of four large ST datasets from platforms like SlideSeq V2, 10X Visium, and MERFISH using these three methods not only reveals exciting biological insights but also demonstrates SMASH’s capability of detecting SVGs that will be otherwise missed by either of the other two methods. A Python-based software implementation of SMASH is available at, https://github.com/sealx017/SMASH-package, which returns the lists of SVGs detected by both SMASH and SPARK-X, allowing users to investigate the overlap between them.

## Results

### Simulations

We evaluated the performance of SMASH, SPARK, and SpaGene in three different simulation setups. We omitted SpatialDE and SPARK from the power comparison for two reasons: a) high computational requirements and b) these two methods have already been greatly studied in previous works [38, 42]. In simulation setup (1), we followed the procedure described in the SPARK-X manuscript [38]. In setups (2) and (3), we considered the Gaussian process (GP)-based spatial regression model from the SpatialDE manuscript [34], respectively with the Gaussian and cosine kernel-based covariance functions (see Equation (2)). In all the setups, three values of the number of cells (*N*) were considered, *N* = 1000, 5000, and 10,000. The spatial coordinates of the cells were simulated first and then the expression levels of *K* (500 or 1000) genes with varying levels of dependence. In setup (1), the expression levels were simulated using a negative binomial distribution, while in setups (2) and (3), those are simulated using a multivariate normal distribution. In all the setups, distinct spatial patterns were ensured to be present in the expression levels. Further details regarding the simulation setups are provided at the end of the Methods section. Figures 1, 2, and 3 respectively correspond to the three simulation setups, in which we display the simulated spatial patterns and the statistical power of the three methods for different parameter combinations.

**Figure 1.**
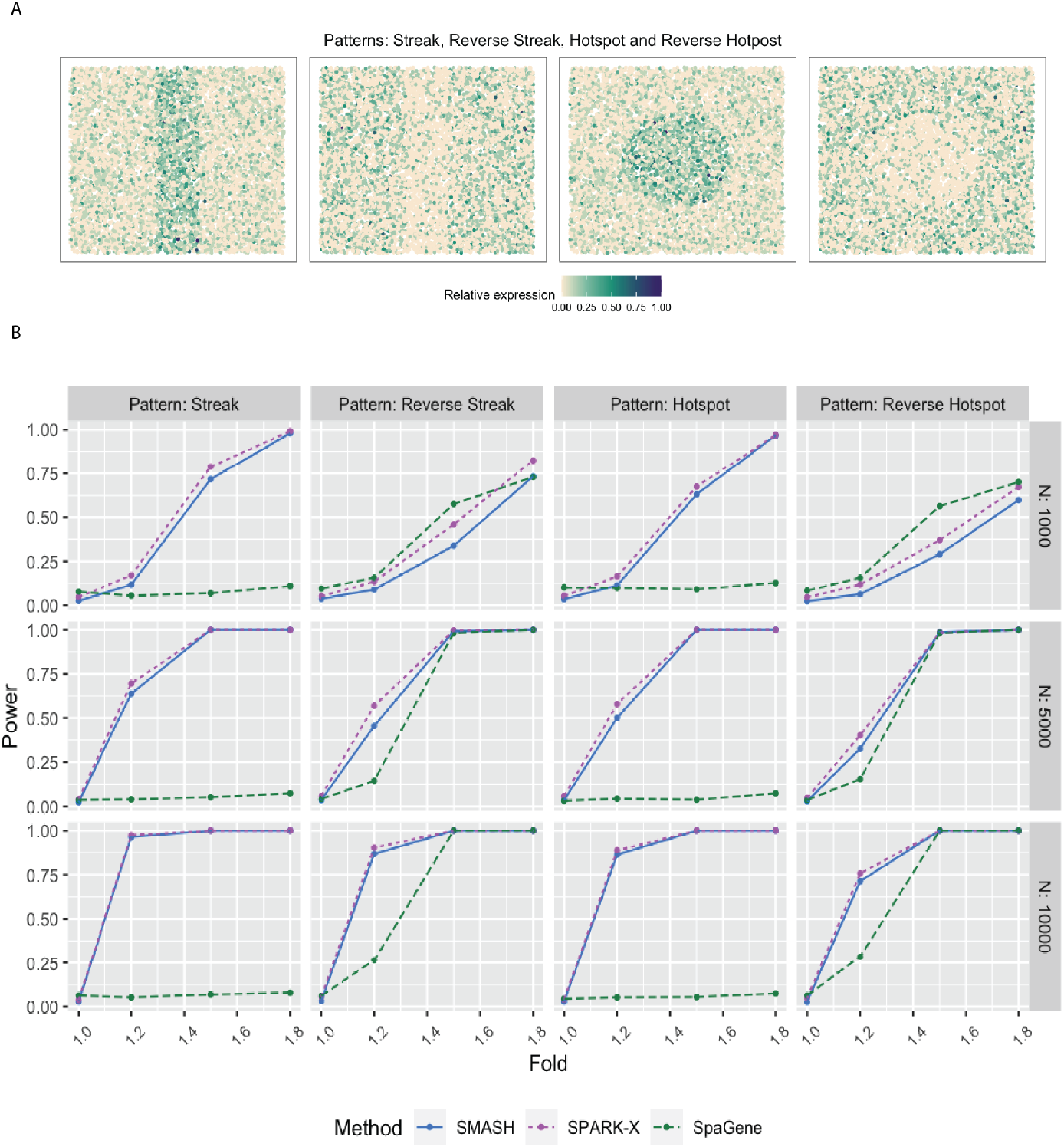
Simulation following the SPARK-X manuscript. **A** Four spatial expression patterns that the genes were assumed to follow. **B** Statistical power plots of the three methods, SMASH, SPARK-X, and SpaGene under varying values of *N* and fold-size, for *K* = 500 genes at a level of *α* = 0.05. The results were averaged over five replications.

**Figure 2.**
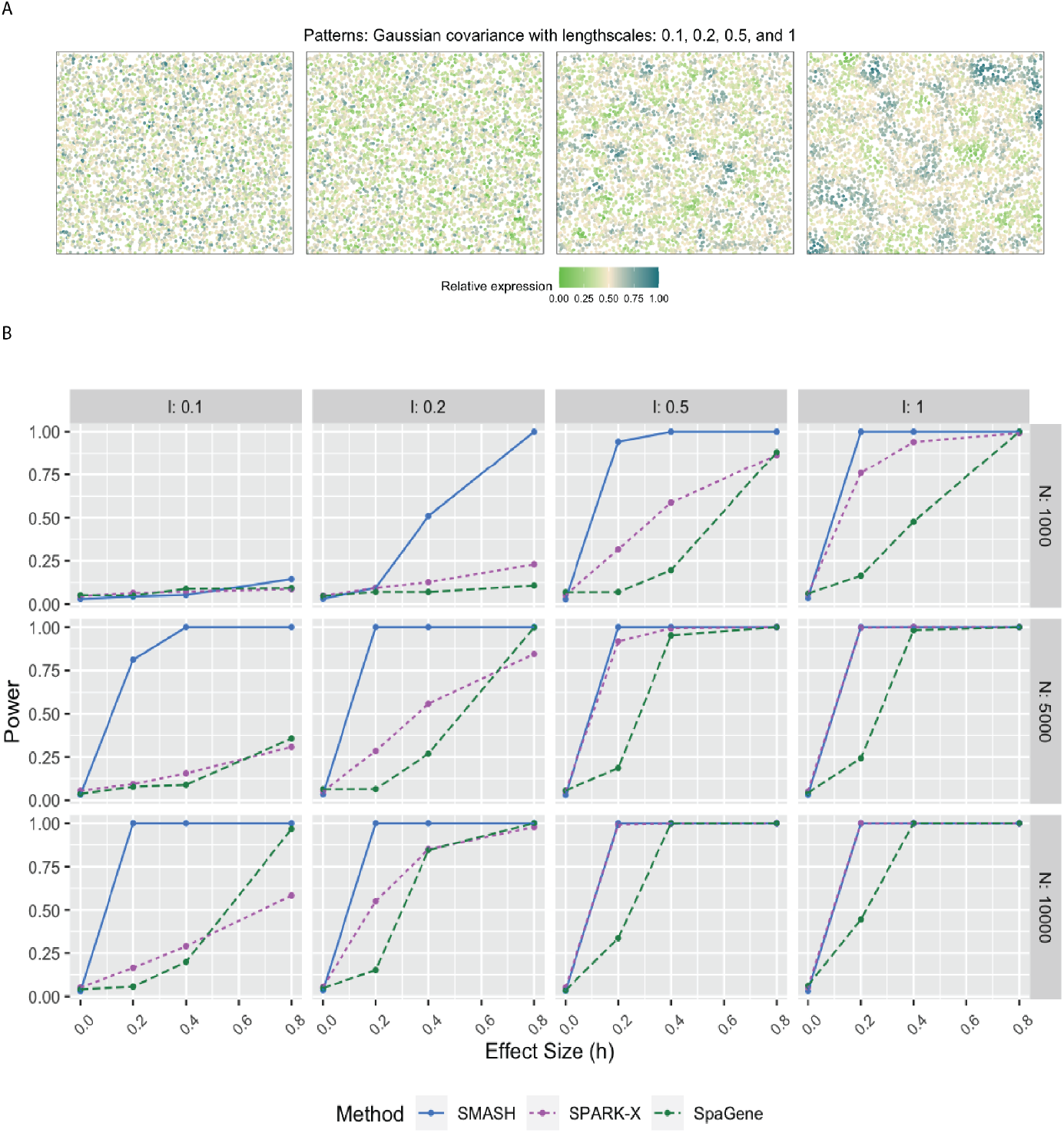
Simulation using Gaussian process-based regression model with the Gaussian covariance. A) Four spatial expression patterns that were generated using Gaussian covariance matrices with four different values of the lengthscale *l*. B) Statistical power plots of the three methods under varying values of *N* and effect-size (*h*) for *K* = 1000 genes at a level of *α* = 0.05. The results were averaged over five replications.

**Figure 3.**
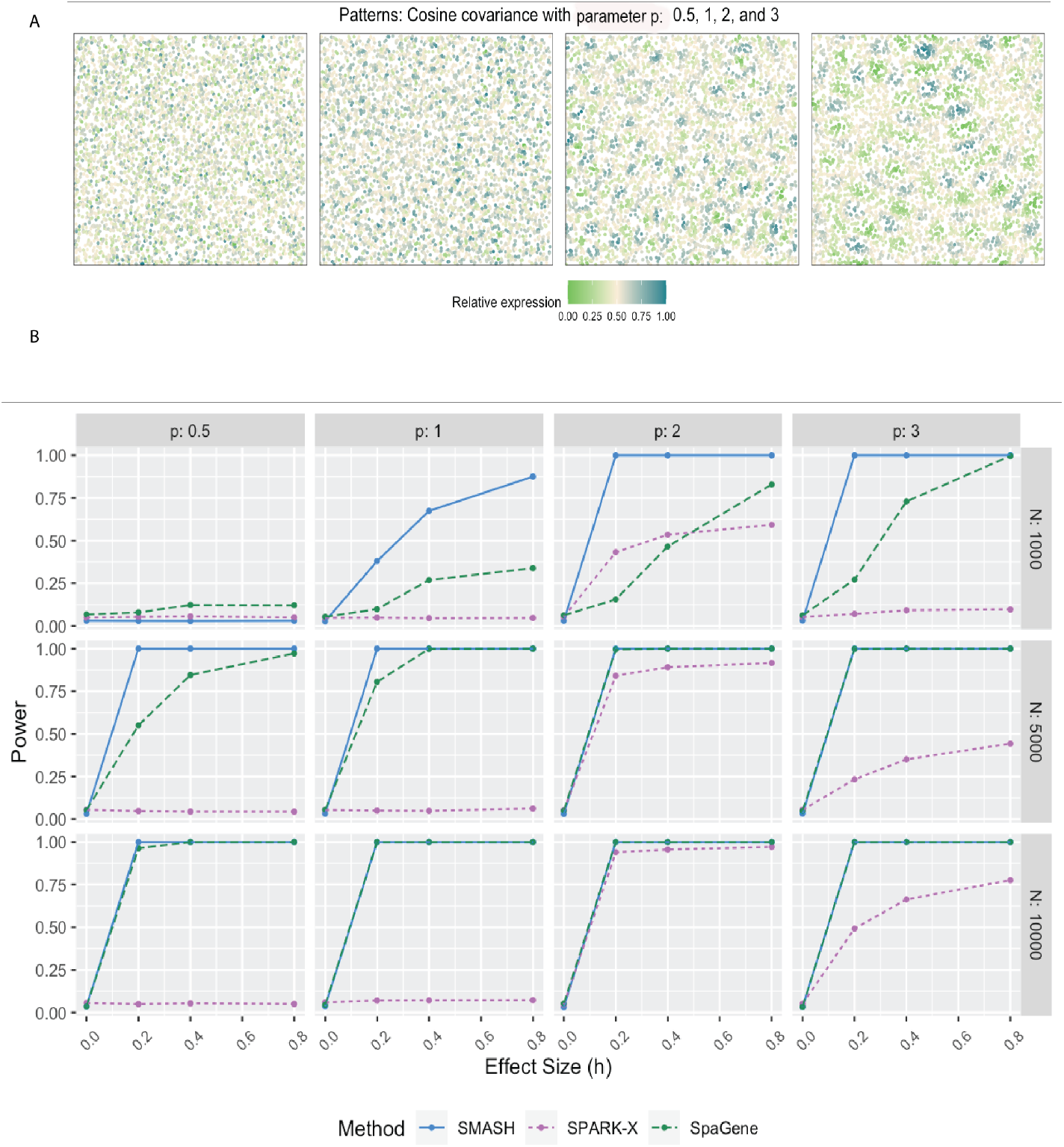
Simulation using Gaussian process-based regression model with the cosine covariance. A) Four spatial expression patterns that were generated using cosine covariance matrices with four different values of the period *p*. B) Statistical power plots of the three methods under varying values of *N* and effect-size (*h*) for *K* = 1000 genes at a level of *α* = 0.05. The results were averaged over five replications.

In simulation setup (1), SMASH, and SPARK-X performed much better than SpaGene for all four spatial patterns, namely streak, reverse streak, hotspot, and reverse hotspot (Figure 1). SpaGene was particularly poor for the patterns: streak and hotspot. The power of SMASH and SPARK-X steadily increased as *N* and the fold-change parameter increased. Note that a fold value of 1 stood for no spatial association while a larger value indicated higher spatial association. This particular simulation setup favored SPARK-X in the sense that the spatial variability of the expression was of the first order, manifesting entirely through the mean or expectation. Even then SMASH could achieve similar power.

In simulation setups (2) and (3), the spatial variability of the expression was of higher order, manifesting through the covariance. In setup (2), which had the Gaussian covariance function, SMASH performed the best followed by SPARK-X and then SpaGene in most cases. SMASH performed the best in setup (3) as well. However, SpaGene achieved better power than SPARK-X here. SPARK-X had almost zero power in many of the cases, especially when the period *p* was small (*p* = 0.5, 1), demonstrating its lack of robustness under complicated spatial dependency structures.

We compared the run-time of the methods in the simulation setup (2) for varying numbers of cells, *N* = 1000, 5000, and 10000 (Table 1). Since the computational complexity of the algorithms mainly differ w.r.t. *N* and not the number of genes *K*, we kept *K* = 1000. We noticed that the run-time of SMASH expectedly increased in an almost squared order w.r.t. *N*. SPARK-X and SpaGene were both extremely fast for just having linear complexity w.r.t. *N*. We also added SpatialDE to this comparison just to show how computationally intensive it can be to fit a fully parametric model in such a context. We omitted SPARK entirely as it is much slower than even SpatialDE with a computational complexity of *O*(*N* ^3^*K*).

**Table 1.** **Computational complexity and run-time comparison. The table lists the theoretical complexity and run-time (in seconds) of the four methods, SMASH, SPARK-X, SpaGene and SpatialDE in a simulation setup with *K* = 1000 genes and varying number of cells *N*. The number of spatial co-ordinates *d* was equal to 2**. ^*∗*^ **SpaGene constructs multiple kNN graphs and performs permutation tests. We are only listing the complexity of the KNN algorithm**.

| Method | Complexity | N = 1000 | N = 5000 | N = 10000 |
| --- | --- | --- | --- | --- |
| SMASH | $O(N^2K)$ | 0.9 | 18.5 | 97.5 |
| SPARK-X | $O(NKd^2)$ | 0.72 | 4.1 | 4.3 |
| SpaGene | $O(NKd)^*$ | 0.2 | 1.1 | 2.1 |
| SpatialDE | $O(N^3 + N^2K)$ | 24.5 | 245 | 971.7 |

### Application to real data

We applied the methods, SMASH, SPARK-X, and SpaGene to four datasets: 1) mouse cerebellum data collected using Slide-seq V2 [21], 2) human dorsolateral prefrontal cortex (DLPFC) data collected using 10X Visium [12], 3) small cell ovarian carcinoma of the ovary hypercalcemic type (SCCOHT) data collected using 10X Visium [12], and 4) mouse hypothalamus data collected using MERFISH [52]. The datasets have varying numbers of genes and spots/cells.

#### Mouse cerebellum by Slide-seqV2

The mouse cerebellum data [21] has 20,117 genes and 11,626 spots. We restricted our focus to the 7,653 genes that express in more than 1% of the spots. The mouse cerebellum is made of four spatial layers, white matter layer (WML), granule layer (GL), Purkinje layer (PL), and molecular layer (ML) [53]. These layers consist of different types of cells. For example, WML contains oligodendrocytes, GL contains granule cells, PL contains Purkinje neurons and Bergmann gila, and ML contains intra-neurons MLI. These cell types can be inferred based on just the transcriptional profiles using cell clustering software like RCTD [32]. We display the inferred cell types overlayed on the spatial locations in Figure 4. Out of the 7,653 genes, SMASH identified 1173 genes to be spatially variable (adjusted *p*-value: *p*_adjust_ *<* 0.05). SPARK-X and SpaGene respectively detected 608 and 518 genes, and the overlaps between the detected SVGs by the three methods are displayed in a Venn diagram (Figure 4). We noted that SPARK-X and SpaGene had many of the SVGs uncommon. SMASH, on the other hand, could identify almost all the detected genes by those two methods, especially SPARK-X, while detecting an additional 363 SVGs.

**Figure 4.**
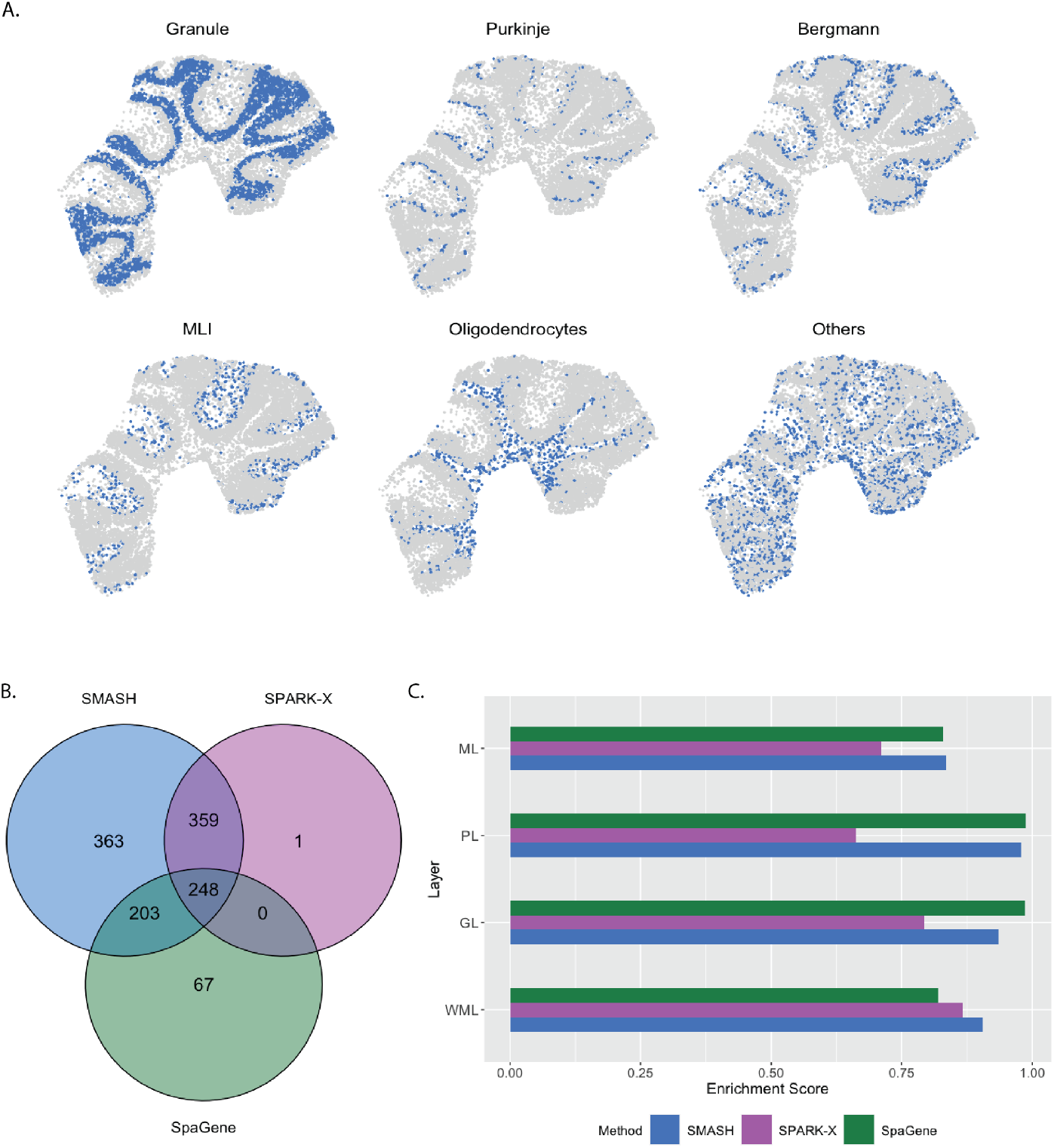
Analysis of mouse cerebellum data. A) Location of the major cell types corresponding to the four spatial layers of the mouse cerebellum. B) Overlap between the detected SVGs by the three methods. C) Enrichment scores of the methods in the four spatial layers.

Next, we performed two types of enrichment analysis. First, we compared the performance of the methods in different layers by computing their enrichment scores (ES) following Liu et al. (2022) [42]. It is based on the expectation that the genes which abundantly express themselves in the four spatial layers, should be identified and ranked top by the methods. In that regard, we noticed that SPARK-X performed poorly in the PL, whereas SpaGene performed poorly in the WML. SMASH, on the other hand, consistently achieved similar or better performance compared to the other two methods in all four layers. Next, we individually performed functional enrichment analysis on the following four sets of SVGs: a) the common genes identified by all three methods, b) the genes identified by SMASH and SpaGene but not by SPARK-X, c) the genes identified by SMASH and SPARK-X but not by SpaGene, and d) the genes identified only by SMASH. The expression pattern of three representative genes of the enriched pathways for each of these four sets of genes, are shown in Figure 5. For set (a), top enriched Gene Ontology (GO) terms, such as GO: 0098916 (anterograde trans-synaptic signaling), GO: 0007268 (chemical synaptic transmission), and GO: 0099536 (synaptic signaling), were broadly associated with synaptic regulation. The protein-coding genes Fam107a, Ppp3ca, and Calm1 appeared in these top pathways. Fam107a seems to express in the PL, whereas the other two express in the GL (Figure 5). For set (b), the top GO terms including GO: 0006873 (intracellular monoatomic ion homeostasis), GO: 0030003 (intracellular monoatomic cation homeostasis), and GO: 0098771 (inorganic ion homeostasis) were associated with ion homeostasis. The representative genes Atp1a3 and Thy1 express in the PL while Calm3 expresses in the GL. For set (c), the top pathways including GO: 0006811 (monoatomic ion transport), GO: 0006812 (monoatomic cation transport), and GO: 0098655 (monoatomic cation transmembrane transport) were associated with ion transportation. The representative genes Pllp and Efnb3 express in the WML, whereas Cox7a2 expresses roughly in the GL. For set (d), the top enriched GO terms, such as GO: 0044057 (regulation of system process) and GO: 0050877 (nervous system process), were associated with regulating different types of system processes. The representative genes Gls, Tmem36a, and Coro2b roughly express in the GL.

**Figure 5.**
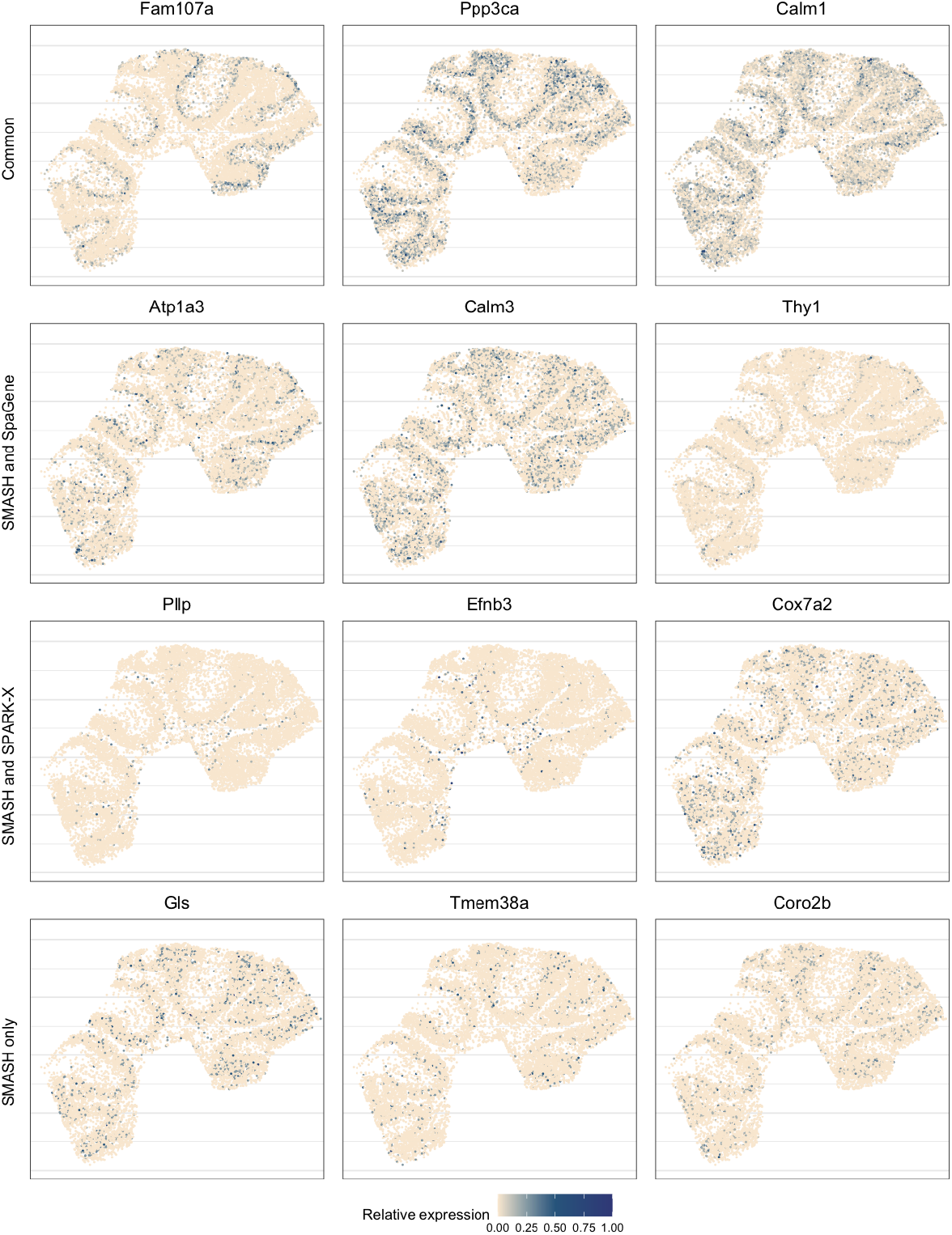
Expression patterns in mouse cerebellum data. Three representative genes from the detected pathways for the four sets of genes: a) the common genes identified by all three methods, b) the genes identified by SMASH and SpaGene but not by SPARK-X, c) the genes identified by SMASH and SPARK-X but not by SpaGene, and d) the genes identified only by SMASH.

#### Human DLPFC by 10X Visium

The human dorsolateral prefrontal cortex (DLPFC) data [12] has 33,538 and 3,639 spots. We focused on the 13,783 genes which express in more than 1% of the spots. Every spot belongs to one of the six manually labeled cortical layers or the white matter layer (WML) (Figure 6). SMASH and SPARK-X identified 10,871 and 10,416 SVGs respectively (*p*_adjust_ *<* 0.05), whereas SpaGene identified only 2379. The overlaps between the detected SVGs by the three methods are displayed in a Venn diagram (Figure 6). We noted that almost all the genes detected by SpaGene were also detected by both SMASH and SPARK-X. SMASH and SPARK-X detected a lot of additional SVGs. We performed functional enrichment analysis on the two sets of detected genes: a) the common genes identified by all three methods and b) the genes identified only by SMASH and SPARK-X but not by SpaGene. For set (a), top enriched GO terms, such as GO: 0099537 (trans-synaptic signaling) and GO: 0099177 (regulation of trans-synaptic signaling), were associated with synaptic signaling. For set (b), top enriched GO terms like GO: 0006397 (mRNA processing) and GO: 0000375 (RNA splicing, via transesterification reactions), were associated with RNA processing. The expression of three representative genes from the set (b) are displayed in Figure 6. There seemed to be a gradient spatial pattern of expression for all three genes which SpaGene failed to detect. Similar to the previous section, we computed the enrichment score (ES) of every method in the seven manually labeled spatial layers. From Figure 6, we noticed that SpaGene performed poorly in terms of ES, especially in Layers 1 and 6. We also performed an additional check as follows. There are three cortical-layer associated SVGs, MOBP, SNAP25, and PCP4, and three blood and immune-related SVGs, HBB, IGKC, and NPY, known to be spatially variable from previous studies [12]. We checked how many of these genes appeared in the lists of the top thousand SVGs (in terms of *p*_adjust_) by the three methods. SMASH and SpaGene respectively ranked five and six of these SVGs, whereas SPARK-X ranked only two cortical-layer associated genes.

**Figure 6.**
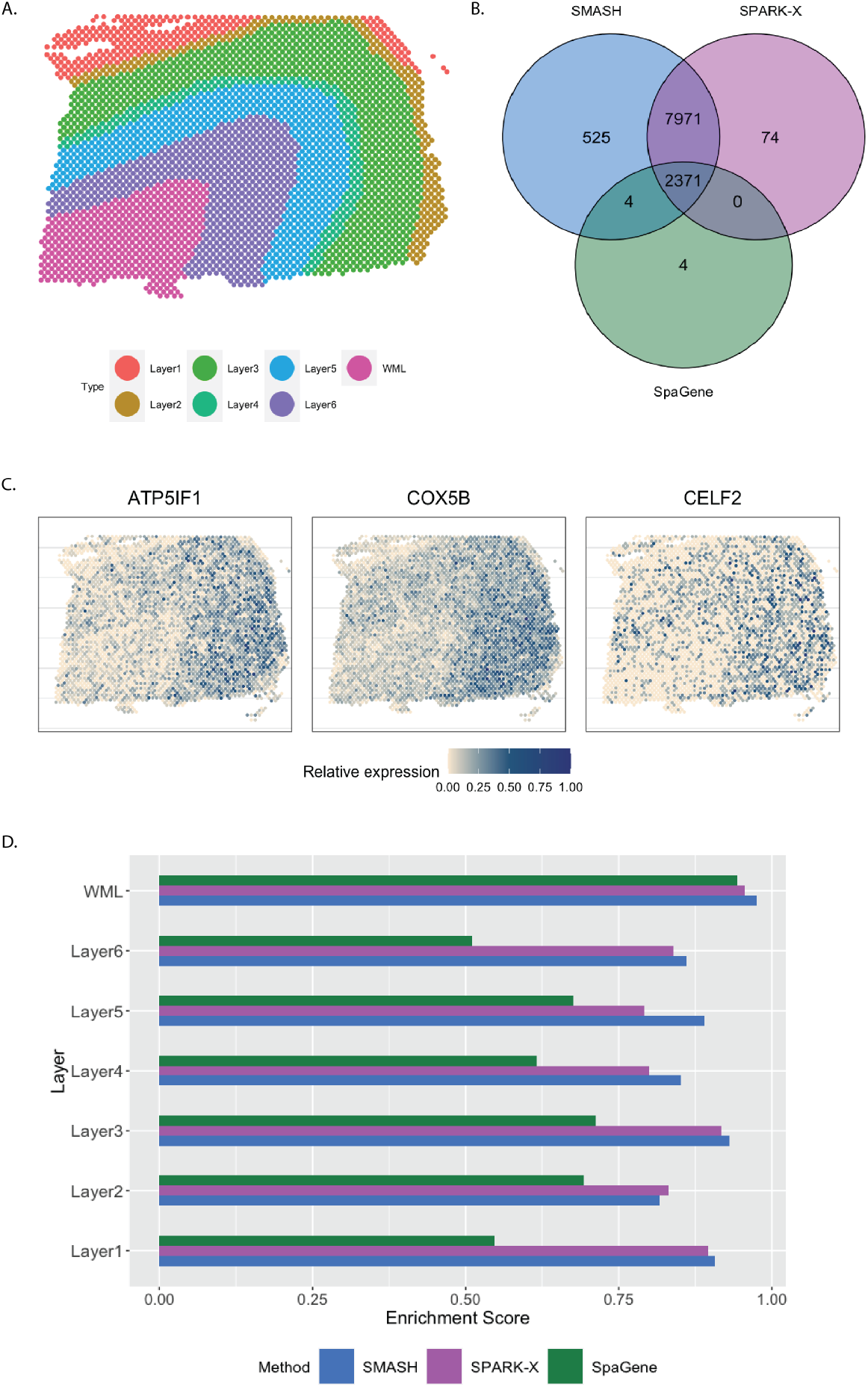
Analysis of human DLPFC data. A) Manually labeled cortical layers (layers 1-6) and white matter layer (WML). B) Overlap between the detected SVGs by the three methods. C) Expression of three representative genes identified only by SMASH and SPARK-X. D) Enrichment scores of the methods in different layers.

#### SCCOHT by 10X Visium

The small cell carcinoma of the ovary hypercalcemic type (SCCOHT) data [54] has 15,229 genes and 2071 cells. We restricted our focus to the 12,001 genes that express in more than 5% of the cells. Sanders et al. (2022) [54] grouped the cells into twelve clusters based on the expression profile of a selected few genes, using Seurat [33], which we display in Figure 7. SMASH, SPARK-X, and SpaGene respectively detected 9361, 6564, and 6899 SVGs (*p*_adjust_ *<* 0.05). The overlaps between the detected SVGs by the three methods are displayed in a Venn diagram (Figure 7). SMASH could detect most of the SVGs identified by at least one of the other two methods and an additional 1634 genes. Similar to the analysis of the Mouse cerebellum data, we checked if the methods could identify the top genes that show enriched expression in the twelve spatially well-separated clusters found by Sanders et al. (2022). We computed the enrichment scores (ES) of the methods for each of the clusters (Figure 7). SMASH achieved consistently higher ES for all the clusters while SpaGene was the second best in most cases. Additionally, in Figure 8, we show the expression of three chosen genes from each of the following four sets of SVGs, a) the common genes identified by all three methods, b) the genes identified by SMASH and SpaGene but not by SPARK-X, c) the genes identified by SMASH and SPARK-X but not by SpaGene, and d) the genes identified only by SMASH. We also checked the clinical relevance of these genes in the existing literature. For example, CITED4, which was detected to be an SVG by all three methods, has been found to be associated with lung adenocarcinoma [55]. From the set (b), ELF4A1 has been found to be associated with gastric cancer [56]. EZH2, from the set (c), is a well-known marker for being associated with the development and progression of different types of cancer [57, 58]. Sanders et al. (2022) [54] also found the expression of EZH2 to be highly variable across their identified spatial clusters. Finally, from the set (d), SEMA4F has been found to be associated with endometrial cancer [59].

**Figure 7.**
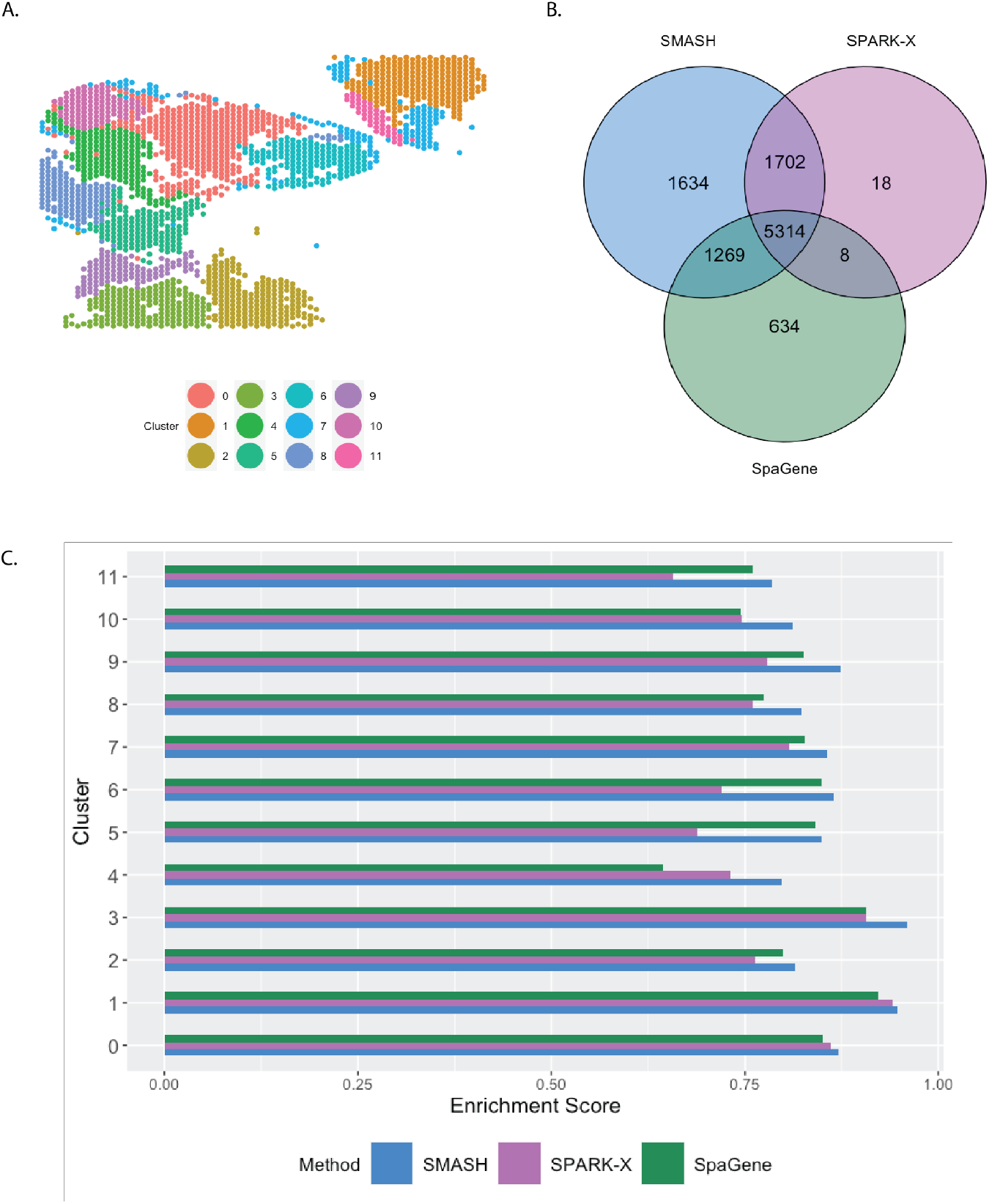
Analysis of SCCOHT data. A) Pre-identified clusters of cells using Seurat. B) Overlap between the detected SVGs by the three methods. C) Enrichment scores of the methods in different clusters.

**Figure 8.**
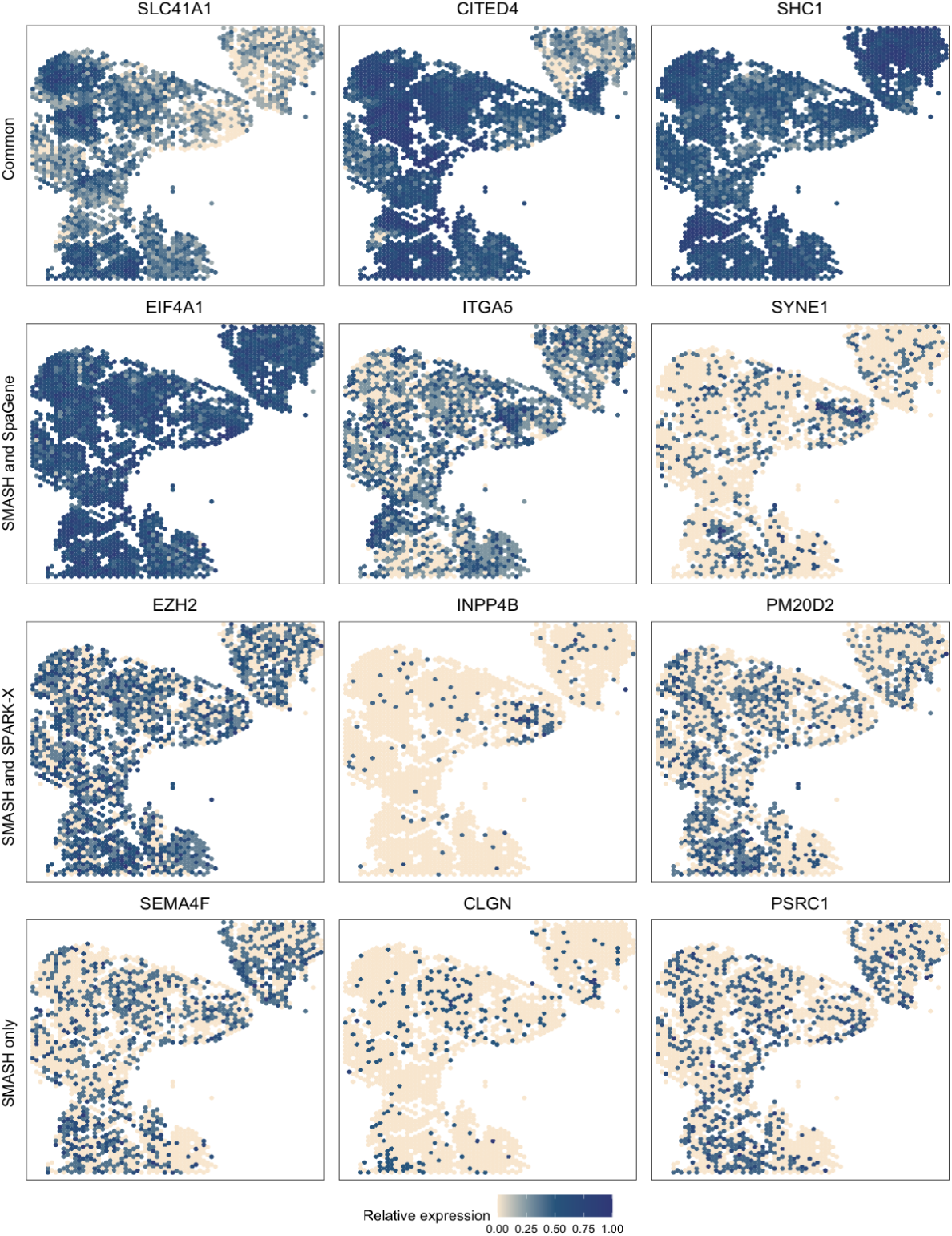
Expression patterns in SCCOHT data. Three representative genes from the four sets of SVGs: a) the common genes identified by all three methods, b) the genes identified by SMASH and SpaGene but not by SPARK-X, c) the genes identified by SMASH and SPARK-X but not by SpaGene, and d) the genes identified only by SMASH.

#### Mouse hypothalamus by MERFISH

The mouse hypothalamus data [52] has 161 genes and 5665 cells. 156 genes are pre-selected markers for different cell types and can thus be expected to be highly variable, whereas the other five are control genes. The cell types, such as endothelial, ependymal, and inhibitory, can be identified based on the transcriptional profiles of the markers. The spatial organizations of a few major cell types are shown in

Figure 9. SMASH was able to detect 139 genes, whereas SPARK-X and SpaGene detected 127 and 124 genes, respectively (*p*_adjust_ *<* 0.01). The overlaps between the SVGs detected by the three methods are shown in Figure 9. SMASH identified all the SVGs SPARK-X could detect, while SpaGene identified one additional SVG. It should be highlighted that all the methods assigned the five control genes to not be spatially variable. We display the expression of two representative genes from three sets of genes, a) the genes identified only by SMASH and SpaGene, b) the genes identified only by SMASH and SPARK-X, and c) the genes identified only by SMASH. We did not focus on the common genes because they have been extensively studied in earlier literature, such as the work of Liu. et al. (2022)[42]. The genes Npy1r and Cplx3 belonged to set (a), and are known to be enriched in inhibitory and excitatory neurons [60, 61]. Rxfp1 and Ntsr1 belonged to set b). Even though both genes are known to express in inhibitory and excitatory neurons, Rxfp1 seems to express in ependymal cells as well. Galr2 and Crhr1 are two genes from set c) which express in multiple cell types including inhibitory cells and astrocytes.

## Discussion

We have proposed a novel non-parametric method SMASH for detecting spatially variable genes (SVGs) in the context of large-scale spatial transcriptomics (ST) datasets. In comparison to existing scalable approaches, SMASH achieves superior power in both complex simulation scenarios and real data analyses while remaining computationally tractable.

**Figure 9.**
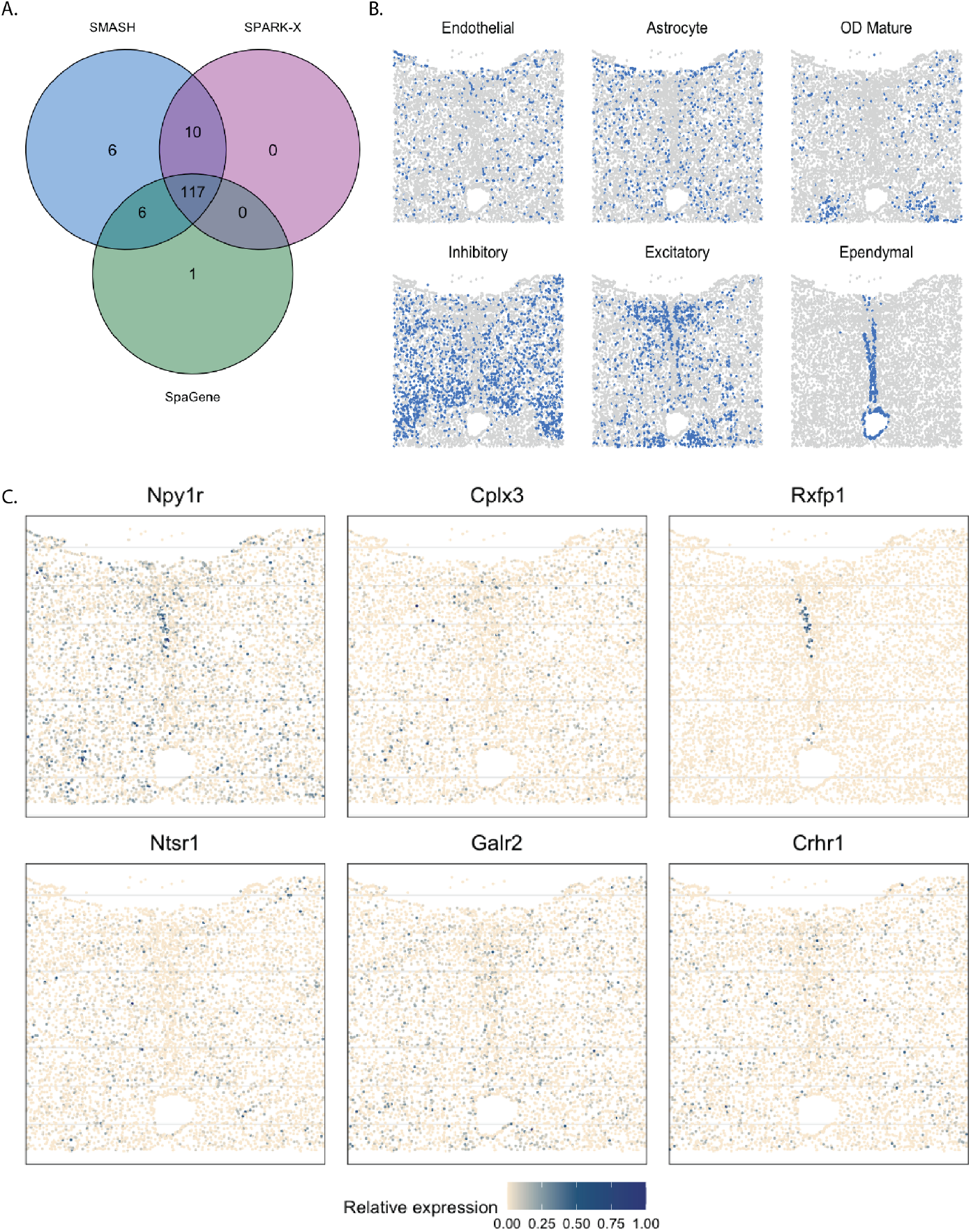
Analysis of mouse hypothalamus data. A) Overlap between the detected SVGs by the three methods. B) Spatial organization of a few major cell types. C) Expression of two representative genes from each of the three sets, a) the genes identified only by SMASH and SpaGene, b) the genes identified only by SMASH and SPARK-X, and c) the genes identified only by SMASH.

Recently developed spatial transcriptomics platforms produce high-dimensional datasets [20, 21, 23] in terms of the number of cells and the number of genes. In such large datasets, fully parametric approaches for detecting SVGs, such as SpatialDE [34] and SPARK [37], albeit statistically powerful, become unusable for their high computational demand. Computationally efficient alternative non-parametric approaches, such as SPARK-X [38] and SpaGene [35], on the other hand, can often turn out to be significantly less powerful. In our method SMASH, we strive to find a balance between these two issues, meaning that we achieve higher statistical power while achieving a moderate degree of scalability. We augment the kernel-based co-variance testing framework [48], used before in SPARK-X, by accounting for more complex spatial dependencies.

In three different simulation setups, one following the SPARK-X manuscript [38] and the other two following the framework of SpatialDE [34], we evaluated the performance of SMASH, along with two other methods: SPARK-X and SpaGene, in terms of type 1 error and power. SMASH achieved consistently similar or better power than the other two methods in all the simulation setups for all combinations of the varying parameters. In contrast, both SPARK-X and SpaGene behaved unpredictably, achieving almost zero detection power in many of the cases. It demonstrated their lack of statistical robustness and failure to capture complicated structures of spatial dependency in the gene expression. In the run-time comparison of the methods, we showed that SMASH, although slower than SPARK-X and SpaGene, remained fairly tractable and was almost ten times faster than a fully parametric approach like SpatialDE. SMASH, SPARK-X, and SpaGene were then applied to four real datasets: 1) mouse cerebellum data collected using Slide-seq V2 [21], 2) human dorsolateral prefrontal cortex data collected using 10X Visium [12], 3) small cell ovarian carcinoma of the ovary hypercalcemic type data collected using 10X Visium [12], and 4) mouse hypothalamus data collected using MERFISH [52]. We compared the methods via a number of avenues: a) checking the overlap between the detected SVGs by the three methods, b) computing enrichment scores (ES) of the methods in different spatial layers or cell types identified based on the transcriptional profiles using popular softwares, such as RCTD [32] and Seurat [33], and c) investigating the functional enrichment of the genes that were detected by SMASH but remained undetected by at least one of the other two methods. For all the datasets, SMASH detected more SVGs than the other two methods, which included nearly all of the SVGs detected by SPARK-X. SMASH could also detect most of the SVGs that were identified by SpaGene but not by SPARK-X. For example, in data (1), from the 7,653 genes after quality control, SMASH identified 1173 SVGs which included 607 out of the 608 SVGs SPARK-X could detect. Out of the 518 SVGs detected by SpaGene, only 248 were also detected by SPARK-X, while SMASH detected 451 of them. It is important to highlight that SMASH produced calibrated *p*-values in the null simulations from all of these datasets, lending credibility to these higher numbers of detected SVGs. In the same dataset, SMASH achieved a higher enrichment score (ES) than the other two methods in different pre-identified spatially separated layers or cell types of the mouse cerebellum. A higher ES meant better capability to identify the genes that showed highly variable expression in a particular spatial layer compared to the rest. In the other datasets as well, SMASH consistently achieved better ES in different spatially localized cell types. We also studied the functional properties and clinical significance of the identified SVGs. For example, in data (3), the gene EZH2 was detected to be spatially variable by SMASH and SPARK-X. EZH2 is a known marker for the progression of multiple types of cancers [57, 58]. It has also previously been found to be highly variable in some particular spatially localized cell clusters by the group of researchers who performed the original experiment [54].

In all the methods we have discussed, including SMASH, the biology of a single tissue section from a single subject is explored at a time. It means that if we either have multiple tissue sections from the same subject or from multiple subjects, the methods will have to identify SVGs individually, disregarding the potential of harnessing shared information between and across the subjects. Thus, we would like to extend SMASH in a hierarchical fashion for jointly analyzing more than one tissue section or subject in the future. One more important functionality that we would like to incorporate would be the ability to classify the genes based on their similarity of spatial expression patterns. For example, SpatialDE [34] considers a hierarchical Bayesian mixture model approach that suffers from extremely high computational demand. SpaGene [42] considers a non-negative matrix factorization [62] of the expression data to identify similarly expressed genes. This approach, although computationally feasible, does not take into account the spatial locations directly and can thus be suboptimal in capturing truly spatial patterns. In the future, we would like to study this problem with a deeper focus and pursue methodological development in this area. Finally, we would like to explore the possibility of using SMASH in the context of multiplex immunohistochemistry (mIHC) datasets [63, 64] where the goal is to identify spatially variable cell types and their interaction.

## Methods

We briefly discuss some of the existing methods such as SpatialDE [34], SPARK [37], SPARK-X [38], and SpaGene [35], and then present the proposed method SMASH. Note that we did not compare SMASH to either SpatialDE or SPARK in our Results section except for the time comparison, primarily due to their high computational demand and the fact that these have already been studied in great detail in earlier works. However, it is crucial to talk about their modeling frameworks for the sake of understanding the other methods better. Let us introduce a few relevant notations. Suppose there is a single subject (image) with *N* cells/spots and the expression profile of *K* genes is observed in the cells. For the *i*-th cell, let *s*_*i*_ denote its location i.e., a vector of spatial (two or three-dimensional) coordinates, and *y*_*ki*_ denote the expression of the *k*-th gene in the cell. Let us also define, *y*_*k*_ = (*y*_*k*1_, …, *y*_*kN*_)^*T*^ and *S* = (*s*_1_, …, *s*_*N*_)^*T*^. For the sake of simplicity, we are assuming that there are no additional covariates but in all the methods, except SpaGene, covariates can be readily incorporated.

### A brief overview of existing methods

#### SpatialDE

SpatialDE uses a Gaussian process (GP)-based spatial regression model [43, 65]. It assumes that at every location *s* ∈ S ⊆ ℝ^3^, the expression of the *k*-th gene is a process *y*_*k*_(*s*) of the following form,

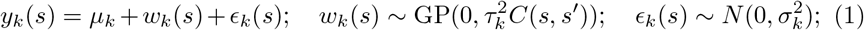

where *w*_*k*_(*s*) is a zero-centered GP with variance 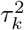 and covariance function *C*(*s, s*′) for *s*′ ∈ 𝒮. A popular choice for covariance function is the Gaussian kernel-based covariance, *C*(*s, s*′) = exp [−||*s* − *s*′||^2^/2*l*^2^], where ||.|| denotes the Euclidean norm, and the hyperparameter *l*, known as the characteristic lengthscale [66], controls the rapidness at which the covariance decays as a function of the spatial distance. *ϵ*_*k*_(*s*) is an independent process with variance 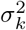. With the observed values of *s* and *y*_*k*_(*s*), the data likelihood corresponding to Equation (1) can then be written as,

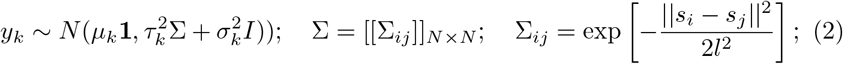

where **1** denotes the *n*-length vector of all 1’s, *I* denotes the *n*-dimensional identity matrix and Σ denotes the corresponding Gaussian covariance matrix. The fixed effect *µ*_*k*_ accounts for the mean expression level and 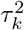 accounts for the expression variance attributable to spatial effects. A large value of 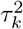 should imply that the gene shows differential spatial expression. To formally test the hypothesis, 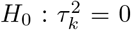 against 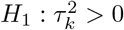, SpatialDE considers the likelihood ratio test (LRT) [67]. To estimate the model parameters under the full model, the log-likelihood corresponding to Equation (2) is optimized w.r.t. (*µ*_*k*_, 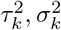) using an efficient algorithm by Lippert et al. (2011) [45]. Ideally, it is desirable to optimize over the hyperparameter *l* as well but for the sake of computational feasibility, *l* is kept fixed at a few carefully chosen values. For every choice of Σ, to analyze all *K* genes, the efficient algorithm requires just one computationally demanding step with a complexity of *O*(*N* ^3^), instead of *O*(*N* ^3^*K*) as incurred in naive algorithms. Along with the Gaussian co-variance function, SpatialDE also considers linear and cosine covariance functions to construct Σ, and finally, combines all the LRT values corresponding to different choices of Σ for the inference. For a particular Σ, the computational complexity of SpatialDE is of *O*(*N* ^3^ + *N* ^2^*K*).

Note that, SpatialDE essentially assumes that gene expression is a continuous random variable (r.v.) and follows a normal distribution. However, for most technologies, gene expression is obtained as count data. Normalizing the count data has been shown to be sub-optimal in several omics studies [68, 69]. Improving upon that, newer methods named SPARK [37] and SPARK-X [38] have been proposed which we discuss next.

#### SPARK and SPARK-X

SPARK [37] considers a generalized linear spatial model (GLSM) [46] with Poisson distribution, which is a generalization of Equation (1) with the following form,

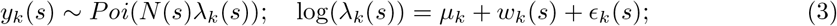

For cell *i, λ*_*k*_(*s*_*i*_) is an unknown Poisson rate parameter that represents the underlying gene expression. The processes *w*_*k*_(*s*) and *ϵ*_*k*_(*s*) are defined as earlier and the variance parameters, 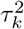 and 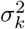 have a similar interpretation as earlier. To test 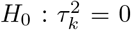, SPARK uses the score test [70]. Parameter estimation and inference are incredibly hard in GLSM which is why SPARK uses an approximate algorithm based on the penalized quasi-likelihood (PQL) approach [47, 71]. The approach has the computational complexity of *O*(*N* ^3^) for every trait, or *O*(*N* ^3^*K*) in total. Thus, it lacks severely in terms of scalability. The SPARK *R* package also has a Gaussian model option (instead of Poisson) which is equivalent to SpatialDE and has a similar computational requirement.

Improving upon SPARK’s scalability, a recent non-parametric method named SPARK-X [38] has been proposed. The method is built on a simple intuition: if *y*_*k*_ is independent of *S*, the spatial distance between two locations *i* and *j* should be independent of the difference in gene expression between the two locations. It computes the expression covariance matrix, 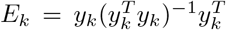 and the distance covariance matrix, *D* = *S*(*S*^*T*^ *S*)^−1^*S*^*T*^ and constructs the test statistic as, 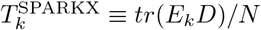 where *tr*() denotes the trace operator. We assume *y*_*k*_ to be mean-standardized here for the sake of simplicity. Under the null hypothesis of no association, *T*_*k*_ asymptotically follows a weighted mixture of independent 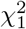 distributions. The weights are the products of the ordered eigenvalues of the matrices, *E*_*k*_, and *D*. This test falls under a general class of covariance tests [48], including the Hilbert-Schmidt independence criteria test [72] and the distance covariance statistic [73]. SPARK-X requires the computational complexity of just *O*(*Nd*^2^) for every gene, or *O*(*NKd*^2^) in total, where *d* is the dimension of the location-space 𝒮, e.g., *d* = 3 if 𝒮 = ℝ^3^. Linearity of the complexity w.r.t. *N* makes SPARK-X easily applicable to large-scale transcriptomics datasets. To capture more complex dependencies between *y*_*k*_ and *S*, SPARK-X also considers several element-wise non-linear transformations of *S* as *g*(*S*), where *g* is a Gaussian or cosine transformation with fixed scale or period parameters. Then, it repeats the above testing procedure replacing *S* with *g*(*S*), and finally, combines all the corresponding *p*-values using Cauchy *p*-value combination rule [74].

However, the form of *D* corresponds to a linear covariance function [75]. It makes SPARK-X equivalent to performing a multiple linear regression of *y*_*k*_ on *S* or *g*(*S*) and testing if the fixed effect parameters differ from zero. Thus, SPARK-X is only capable of detecting first-order spatial dependencies and as shown in the Results section, severely lacks power for higher-order dependencies. Refer to the supplementary information for more discussion on this topic.

#### SpaGene

A very recently developed method, SpaGene [42] is quite different from the rest of the methods in the sense of being model-free and based on graphs. The intuition behind the method is that the cells/spots with high gene expression are more likely to be spatially connected than random. It constructs the *k*-nearest neighbor (kNN) graph based on spatial locations. Then, for each gene, it extracts a subnetwork comprising only cells/spots with high expression from the kNN graph. SpaGene quantifies the connectivity of the subnetwork using the earth mover’s distance (EMD) [50] between degree distributions of the subnetwork and a fully connected one. To generate the null distribution of the EMD for inference, a permutation test is considered. For further details, we refer the readers to [41].

### Proposed method: *SMASH*

We consider a test statistic based on the kernel-based test of independence developed by Zhang et al. (2012) [48] as, 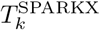 ≡ *tr*(*E*_*k*_*H*(*S*))/*N*, where *E*_*k*_ is defined as earlier and *H*(*S*) is any *N*×*N* kernel-based covariance matrix based on the locations *S*. In this work, we consider *H*(*S*) to have three forms: a) the Gaussian kernel-based covariance matrix, Σ defined in Equation (2), b) a cosine kernel-based covariance matrix of the form, *H*(*S*) = [[cos(2*π*||*s*_*i*_ − *s*_*j*_||*/p*)]]_*N*×*N*_, where parameter *p* is known as the period, and c) the linear kernel-based covariance matrix *D* considered in SPARK-X (Gaussian and cosine transformations of *S*: *g*(*S*) are considered as well in constructing *D*). For the Gaussian covariance matrix, we consider ten data-driven fixed values of the lengthscale *l* [66], and for the cosine covariance matrix, we consider ten data-driven fixed values of the period *p*. For detailed discussions on the relevance of using these types of covariance matrices in the context of ST datasets, we refer the readers to [34] and [37].

Similar to the SPARK-X test statistic, 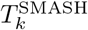, the asymptotic null distribution of 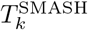 will be a weighted mixture of independent 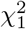 distributions, where the weights are the products of the ordered eigenvalues of the matrices, *E*_*k*_, and *H*(*S*). However, *H*(*S*) does not always have a form like *D* and thus, its eigenvalues can not be computed with the complexity of *O*(*Nd*^2^). Instead, it requires the complexity of *O*(*N*^3^), which becomes intractable as *N* increases. It has been shown that the asymptotic null distribution of 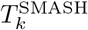 can be well approximated by a gamma distribution [48] as below,

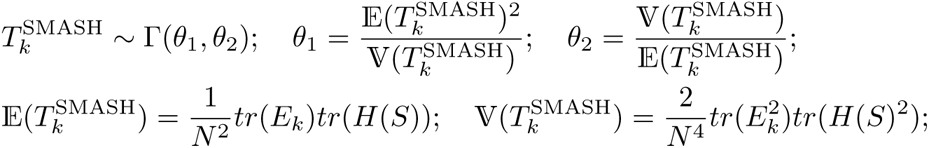

where 𝔼() and 𝕍() denote the expectation and variance, respectively. It is easy to verify that 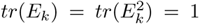. Notice that we can now avoid any operation of complexity *O*(*N* ^3^). Computation of *tr*(*H*(*S*)^2^) just requires the complexity of *O*(*N* ^2^) using the property that 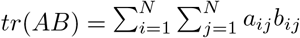 for two matrices, *A* = [[*a*_*ij*_]]_*N*×*N*_ and *B* = [[*b*_*ij*_]]_*N*×*N*_. Thus, for a particular choice of *H*(*S*), to analyze all *K* genes, SMASH requires the complexity of *O*(*N* ^2^*K*). This computational complexity is higher than SPARK-X. But we are making that sacrifice to gain significantly more power, as shown in both simulation studies and real data analyses while still achieving a moderate degree of scalability. It is worth pointing out that even though SMASH is non-parametric and does not make any distributional assumptions, 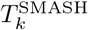 shares a close connection with the SpatialDE model under some additional assumptions (see the supplementary information).

As mentioned earlier, we consider multiple (say, *R*) choices for *H*(*S*), to construct multiple test statistics: 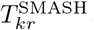, *r* = 1, …, *R*. Finally, we combine the *p*-values corresponding to these test statistics using the minimum *p*-value combination rule [74] (see the supplementary information for more details). Note that we have assumed that *y*_*k*_ is mean-standardized and there are no additional covariates to be taken into account. In presence of covariates, we would regress the covariates out from the gene expression vector *y*_*k*_, prior to performing the test, using a multiple linear regression model. To further elaborate, letting *X* be the corresponding matrix of covariates, we would compute the projection matrix *P*_*X*_ = *X*(*X*^*T*^ *X*)^−1^*X*^*T*^, and substitute the vector *y*_*k*_ with 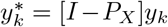, in our proposed test statistic. A Python-based software implementation of SMASH is available at, https://github.com/sealx017/SMASH-package. The software returns the SVGs detected by both SMASH and SPARK-X giving users a chance to examine the overlap between the methods.

### FDR control

In the real data analysis, we used Benjamini-Yekutieli [76] procedure to control the false discovery rate (FDR) at 0.05 (or, 0.01) for all the methods. In the Results section, *p*_adjust_ refers to the adjusted *p*-values. It was shown by Zhu et al. (2021) [38] that parametric methods like SpatialDE and SPARK often produce highly inflated *p*-values for most ST datasets, and hence need additional testing correction. To check if our *p*-values were inflated in the four real datasets, we randomly permuted the spatial locations of the cells/spots five times and then performed the tests using the three methods. Thus, we obtained the empirical null distribution of the *p*-values for each method which we displayed as quantile-quantile plots (see Figure S1 in the supplementary information). In all four cases, SMASH showed no sign of inflation with rather slightly conservative *p*-values which is expected since the minimum *p*-value combination rule used for combining the *p*-values in our method, is known to be conservative [77].

### Enrichment scores

In the real data analysis, we computed the enrichment scores (ES) of the three methods following the procedure outlined in Liu et al. (2022) [42]. Cell clustering based on biological knowledge or using popular softwares, such as RCTD [32] and Seurat [33], with the transcriptional profiles, can often identify spatially localized layers or cell types. Therefore, marker genes in those spatially-restricted cell types should ideally be identified as SVGs. Suppose there are *M* cell types. For every cell type *m*, the gene set *G*_*m*_ is built from the top 50 markers based on the fold change between the expression in the cell type *m* compared to the others. The SVGs detected by the three methods are ranked from the most to the least significant. Finally, unweighted gene set enrichment analysis [78] is implemented to evaluate the enrichment of the gene sets, *G*_*m*_, *m* = 1, …, *M*, in the high ranking of the ranked SVG lists of the methods.

### Simulation description

In simulation setup (1), we generated the spatial coordinates for varying numbers of cells, *N* = 1000, 5000, and 10,000 using a random point-pattern Poisson process [79]. The expression values of *K* = 500 genes in these cells were simulated based on a negative binomial distribution displaying one of the four spatial patterns: streak, reverse streak, hotspot, and reverse hotspot as shown in Figure 1. For each of the patterns, 80% of the spatial locations were assumed to be background locations, while the rest 20% were assumed to be part of the pattern. The difference between the mean expression of a gene on a background location and a patterned location was captured through a fold-change parameter. Several values of fold-change were considered where a value of 1 implied a null scenario i.e., no spatial pattern, and a high value implied a prominent spatial pattern. We refer to Zhu et al. (2021) [38] for more details.

For simulation setup (2), we considered the Gaussian process (GP)-based spatial regression model from SpatialDE [34]. The locations were simulated based on Uniform distribution, which were then used to construct Gaussian covariance matrices with varying lengthscale (*l*) parameters as in Equation (2). The expression levels of genes were independently and identically simulated from the multivariate normal distribution described in Equation (2) for different values of the variance parameters 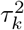 and 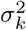. We fixed the total variance, 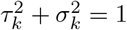, and varied the individual values as 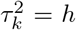 and 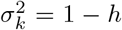, where “effect-size” *h* ranged from zero to larger values implyinh null to an increasingly stronger spatial pattern. In simulation setup (3), we followed setup (2) replacing the Gaussian covariance with the cosine covariance for varying values of the period parameter *p*. In all three setups, we compared SMASH, SPARK-X, and SpaGene in terms of type 1 error and power.

## Additional Files

**Additional file 1: Supplementary information**. It includes additional mathematical discussions pertaining to the proposed method, and one figure displaying the empirical null *p*-values of the three methods, SMASH, SPARK-X, and SpaGene in the four real datasets in consideration.

## Acknowledgements

We would like to thank Dr. Kristen Wells-Wrasman for her help with processing the SCCOHT dataset.

## Availability of data and materials

A Python-based software implementation of SMASH is available at, https://github.com/sealx017/SMASH-package. The package provides two detailed notebooks to perform the analysis on the mouse hypothalamus data by MERFISH and the human DLPFC data by 10X Visium (along with the datasets as compressed Python objects).

Both the mouse cerebellum data by Slide-seqV2 and the human DLPFC data by 10X Visium are available in the *R* Bioconductor package: STexampleData [80], available at, https://bioconductor.org/packages/release/data/experiment/html/STexampleData.html. The full mouse hypothalamus data by MERFISH is available at, https://datadryad.org/stash/dataset/doi:10.5061/dryad.8t8s248, from which we focused on only “Replicate 6”, as it had the largest number of cells and highest confluency [34]. The SCCOHT dataset by 10X Visium was collected at the University of Colorado Denver - Anschutz Medical Campus, and is available upon request.

