## Supplementary information for "SMASH: Scalable Method for Analyzing Spatial Heterogeneity of genes in spatial transcriptomics data"

### 1 Further details on the SMASH test statistic

#### 1.1 Choice of kernel covariance matrices

In the main text, we define the SMASH test statistic for a gene $k$ to have the following form,

$$T_k^{\text{SMASH}} \equiv \frac{tr(E_k H(S))}{N}; \quad E_k = y_k(y_k^T y_k)^{-1} y_k; \tag{1}$$

where $H(S)$ is any $N \times N$ kernel-based covariance matrix based on the locations $S$ and $y_k$ is the mean-standardized expression vector of the gene $k$. As mentioned in the main text, when additional covariates are present, we would replace $y_k$ by $y_k^* = [I - P_X]y_k; P_X = X(X^T X)^{-1} X^T$, where $X$ is the design matrix. We consider $H(S)$ to have following three forms:

1. The Gaussian kernel-based covariance matrix:

$$H(S) = \left[\left[\exp\left(-\frac{||s_i - s_j||}{2l^2}\right)\right]\right]_{N\times N}; \quad ||s_i - s_j||= \sqrt{(s_{i1} - s_{j1})^2 + (s_{i2} - s_{j2})^2}$$

for ten values of the lengthscale parameter $l$.

2. The cosine kernel-based covariance matrix of the form:

$$H(S) = \left[\left[\cos\left(\frac{2\pi||s_i - s_j||}{p}\right)\right]\right]_{N\times N}$$

for ten values of the period parameter $p$.

3. The linear kernel-based covariance matrix of the form:

$$H(S) = g(S)(g(S)^T g(S))^{-1} g(S)^T$$

where three choices of the coordinate-wise transformation $g$ are considered:

- An identity transformation i.e., $g(s_{i1}) = s_{i1}, g(s_{i2}) = s_{i2}$ for $i = 1, \ldots, N$, or, $g(S) = S$.

- A Gaussian transformation as $g(s_{i1}) = \exp(-s_{i1}^2/2t_1^2), g(s_{i2}) = \exp(-s_{i12}^2/2t_2^2)$ for $i = 1, \ldots, N$, for five values of scale parameters $t_1$ and $t_2$.

- A cosine transformation as $g(s_{i1}) = \cos(2\pi s_{i11}/\phi_1), g(s_{i2}) = \cos(2\pi s_{i12}/\phi_2)$ for $i = 1, \ldots, N$, for five values of period parameters $\phi_1$ and $\phi_2$.

The Gaussian and cosine kernel-based covariance matrices have earlier been used in SpatialDE [5] and SPARK [4]. We follow SpatialDE to choose a set of fixed grid points for both the lengthscale parameter $l$ and the period $p$. In particular, we first obtain the minimum ($d_{min}$) and the maximum ($d_{max}$) values of the non-zero Euclidean distances across all pairs of spatial locations. Then, we extract ten equally-spaced values between $\log_{10}(d_{min}/2)$ and $\log_{10}(d_{max}/2)$. These values are then converted to the original scale by taking the power of ten and subsequently used as the values of $l$ and $p$. Let us denote the test statistics corresponding to these twenty covariance matrices (10 each for the Gaussian and cosine kernels) as, $T_{kr}^{\text{SMASH}}, r = 1, \ldots, 20$. The linear kernel-based covariance matrices with transformed coordinates ($g$-transformation) have been earlier used in SPARK-X [6]. Following SPARK-X, we vary the transformation parameters, $t_1$, $t_2$, $\phi_1$ and $\phi_2$ to be the $20\%, 40\%, 60\%, 80\%$, and $100\%$ quantiles of the absolute values of the $x$ and $y$ coordinates in the data. Let us denote the test statistic corresponding to these eleven (1 for the identity transformation and five each for the Gaussian and cosine transformations) linear kernel-based covariance matrices as, $T_{kr}^{\text{SMASH}}, r = 21, \ldots, 31$. We denote the $p$-values corresponding to each $T_{kr}^{\text{SMASH}}$ as $p_{kr}$. Then, we can get SPARK-X's result by combining the $p$-values, $p_{kr}, r = 21, \ldots, 31$ using a Cauchy combination rule [2]. Note that the same Gaussian and cosine kernel-based covariance matrices ($r = 1, \ldots, 10$) are also used in SpatialDE [5] but not in SPARK-X. The linear kernel-based covariance matrix with identity transformation ($r = 21$) is used in both SpatialDE and SPARK-X, while the other transformations ($r = 22, \ldots, 31$) are uniquely considered in SPARK-X. We can get an approximate version of the SpatialDE (more in Section 1.2) $p$-value by combining $p_{kr}, r = 1, \ldots, 21$ using a simple minimum $p$-value approach as, $p_{k,comb1} = 21 * \min\{p_{k1}, \ldots, p_{k21}\}$. To combine the SPARK-X specific $p$-values ($r = 22, \ldots, 31$), we follow the same Cauchy combination rule [2] to get $p_{k,comb2}$. To construct the final $p$-value of SMASH, we perform a minimum $p$-value combination as, $p_{k,final} = 2 * \min\{p_{k,comb1}, p_{k,comb2}\}$. We do acknowledge that using two levels of minimum $p$-value combination can be conservative in some cases, which is partly observed in the null simulation studies performed using the real datasets in Section 3. However, our focus was on preventing the high degree of inflation in the $p$-values routinely observed in model-based methods like SpatialDE and SPARK.

#### 1.2 Similarity with SpatialDE

From Equation (2) of the main text, recall that SpatialDE considers the following model,

$$y_k \sim N(\mu_k \mathbf{1}, \tau_k^2 \Sigma + \sigma_k^2 I)); \quad \Sigma = [[\Sigma_{ij}]]_{N \times N}; \quad \Sigma_{ij} = \exp\left[-\frac{||s_i - s_j||^2}{2l^2}\right];$$

SpatialDE optimizes two likelihood functions: one corresponding to the 'full' model shown above and the other corresponding to the reduced 'null' model without the $\tau_k^2\Sigma$ component in the covariance. Then, to test the null hypothesis, $H_0 : \tau_k^2 = 0$, it constructs a likelihood ratio test (LRT) comparing the optimized values of the above two likelihood functions. If $y_k$ is mean-standardized i.e., $\mu_k = 0$, the corresponding score test statistic [1] can easily be shown to have the form,

$$\begin{aligned}
T_{score} &= y_k^T \Sigma y_k \\
&= tr(y_k y_k^T \Sigma) \\
&= (y_k^T y_k) tr(E_k \Sigma) \quad \text{since } (y_k^T y_k) \text{ is a scalar and } E_k = y_k(y_k^T y_k)^{-1} y_k, \\
&= N(y_k^T y_k) T_k^{\text{SMASH}} \quad \text{when } H(S) = \Sigma \text{ in Equation (1).}
\end{aligned}$$

Thus, we have derived a rough similarity between the SMASH test statistic with the framework of SpatialDE.

#### 2 SPARK-X's equivalence with a multiple linear regression model

Let us consider the following linear regression model with $y_k$ as the dependent variable and the columns of the spatial location matrix $S$ ($x$ and $y$ coordinates) as the predictors,

$$y_k = S\beta_k + \epsilon_k; \quad \epsilon_k \sim N(0, \sigma_k^2 I)$$

The fixed effect coefficients vector $\beta_k$ is of length 2 and its OLS estimate has the form, $\hat{\beta}_k = (S^T S)^{-1} S^T y_k$ with an estimated covariance matrix of $var(\hat{\beta}_k) = \hat{\sigma}_k^2 (S^T S)^{-1}$, $\hat{\sigma}_k^2 = y_k^T (I - D) y_k / N$; where $D = S(S^T S)^{-1} S^T$ from the main text. To test the null hypothesis, $H_0 : \beta_k = 0$, we would consider the following Wald test statistic,

$$
\begin{aligned}
T &= \hat{\beta}_k^T(var(\hat{\beta}_k))^{-1}\hat{\beta}_k/N \\
&= y_k^T S(S^TS)^{-1}(S^TS)(S^TS)^{-1}S^T y_k/(N\hat{\sigma}_k^2) \\
&= y_k^T S(S^TS)^{-1}S^T y_k/(N\hat{\sigma}_k^2) \\
&= tr((y_k y_k^T)(S(S^TS)^{-1}S^T))/(N\hat{\sigma}_k^2) \\
&= (y_k^T y_k) tr(E_k(S(S^TS)^{-1}S^T))/(N\hat{\sigma}_k^2) \quad \text{since } (y_k^T y_k) \text{ is a scalar and } E_k = y_k(y_k^T y_k)^{-1}y_k^T, \\
&= [(y_k^T y_k)/\hat{\sigma}_k^2][tr(E_k D)/N] \quad \text{where } D = S(S^TS)^{-1}S^T \text{ from the main text,} \\
&= [(y_k^T y_k)/\hat{\sigma}_k^2]T_k^{\text{SPARKX}} \quad \text{where } T_k^{\text{SPARKX}} = tr(E_k D)/N \text{ from the main text.}
\end{aligned}
$$

Thus, we have shown that the SPARK-X test statistic $T_k^{\text{SPARKX}}$ is proportional to the simple Wald test statistic obtained from a multiple linear regression model. However, the asymptotic distributional assumptions under null are different in both cases. Note that SPARK-X also replaces $S$ by $g(S)$ where $g$ is a coordinate-wise transformation function as discussed in Section 1.1.

### 3 QQ-plots under null simulations

As mentioned in the main text, we performed null simulation studies to construct an empirical null distribution of the $p$-values for every method. With each of the four datasets, we randomly permuted the cell/spot coordinates five times and then applied the three methods, SMASH, SPARK-X, and SpaGene to obtain the respective $p$-values. The $p$-values were then transformed as $-\log_{10}(p\text{-value})$ and displayed against the expected values as quantile-quantile plots (QQ-plots) in Figure S1. We noticed that SMASH showed no sign of $p$-value inflation and was rather slightly conservative. It is expected since the minimum $p$-value combination rule we use, is known to be conservative [3] (see Section (1.1)). SpaGene produced slightly inflated $p$-values in the SCCOHT dataset while SPARK-X did not show any sign of inflation.

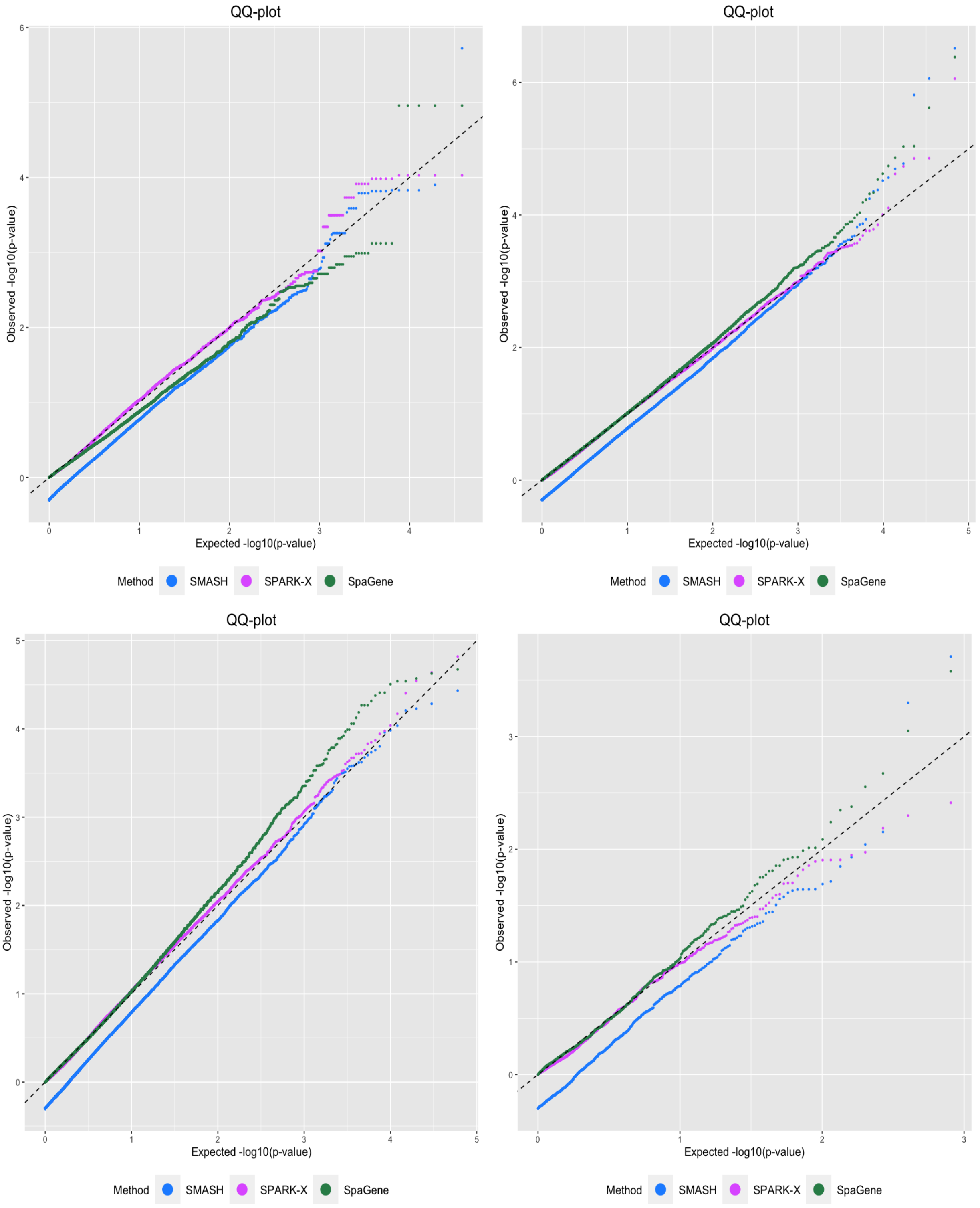


Figure S1: Top left: Mouse cerebellum by Slide-seqV2, top right: Human DLPFC by 10X Visium, bottom left: SCCOHT by 10X Visium, bottom right: Mouse hypothalamus by MERFISH
